# Global Signal Removal (GSR) as graph spatial filtering

**DOI:** 10.64898/2026.04.06.716832

**Authors:** Fahimeh Arab, Benjamin S. Sipes, Srikantan S. Nagarajan, Ashish Raj

**Author notes:** These authors contributed equally to this work (Fahimeh Arab, Benjamin S. Sipes).

## Abstract

Global Signal Removal (GSR) is a widely applied step in functional magnetic resonance imaging (fMRI) preprocessing. Although GSR conventionally denotes ‘Global Signal Regression,’ we use ‘Global Signal Removal’ to encompass a broader family of spatial filtering operations. GSR in general remains controversial due to concerns about introducing spurious anticorrelations and removing neurally meaningful signals. In this paper, we provide a precise geometric characterization by formalizing GSR as graph spatial filtering. We demonstrate that the most common form of GSR, Regression-GSR, equates to a rank-1 deflation of the covariance matrix (i.e. functional connectivity) by the degree vector. Empirically, the degree vector is dominated by the first principal component of the functional connectivity matrix (correlation = 0.88 ± 0.12 in resting-state HCP data), making Regression-GSR an approximation to first eigenmode removal. This view of GSR as a spatial projection framework allows us to develop a family of GSR variants, each expressible in a unified spatial filter: Naive-GSR removes the uniform vector, PCA-GSR precisely removes the first eigenvector, and SC-GSR, a new variant we introduce that removes the first harmonic of the structural connectivity matrix. A key distinction emerges: while Naive, PCA, and SC-GSR are orthogonal projections, Regression-GSR is an oblique projection that computes regional weights proportional to the degree vector but removes a spatially uniform signal. All GSR variants induce numerical singularity in the covariance matrix, but they differ in their effects on task-state separability, which we examine empirically. In summary, we reframe GSR as a family of graph spatial filters that enable interpretability of its effects, with systematically varying effects on network connectivity across variants.

## Introduction

Global Signal Removal (GSR) remains one of the most widely applied yet vigorously debated preprocessing steps in functional MRI (fMRI) analysis. Originally introduced to remove non-neuronal confounds such as respiration and head motion, its validity has been challenged by concerns that the “global signal” contains behaviorally relevant neural information (Aguirre et al., 1998; Liu et al., 2017; Murphy & Fox, 2017). Despite decades of research and comprehensive benchmarking of preprocessing strategies (Parkes et al., 2018), the field has yet to reach a consensus on its use, with some studies demonstrating that GSR enhances the prediction of behavioral traits (Li, Kong, et al., 2019), while others warn that it systematically distorts group comparisons (Saad et al., 2012, 2013), particularly in clinical populations such as Autism Spectrum Disorder (Gotts et al., 2013).

The controversy surrounding GSR has largely centered on its statistical consequences. A seminal critique by Murphy et al. demonstrated that GSR mathematically mandates a zero-sum constraint on the data, inevitably introducing artificial negative correlations (anticorrelations) even in the absence of true biological antagonism (Murphy et al., 2009; Wong et al., 2012). However, anticorrelations between the default mode and task-positive networks have been shown to persist without GSR when the global signal is removed via component-based methods such as CompCor, suggesting they reflect genuine neurobiological antagonism rather than statistical artifact (Behzadi et al., 2007; Chai et al., 2012). The anticorrelations in functional connectivity can alter graph theoretical metrics such as modularity and path length (Gargouri et al., 2018; Saberi et al., 2021), and bias estimates of effective connectivity in dynamic causal models (Almgren et al., 2020).

Conversely, proponents argue that widespread signal deflections driven by motion and physiology are major sources of non-neural global signal variance (Aquino et al., 2020; Bolt et al., 2025; Power, Laumann, et al., 2017; Power, Plitt, et al., 2017), with recent work showing that global brain activity itself can induce bias in fMRI motion estimates (Mao et al., 2024) and that failure to remove them leaves connectivity estimates vulnerable to artifacts. Direct electrophysiological evidence corroborates this view: using simultaneous EEG-fMRI recordings, Xifra-Porxas et al. (2025) demonstrated that systemic physiological fluctuations account for a significantly larger fraction of global signal variance than electrophysiological activity, and that GSR preserves connectivity patterns associated with neural alpha and beta oscillations. Similarly, coupled hemodynamic and cerebrospinal fluid oscillations observed during sleep further support the physiological basis of global signal fluctuations (Fultz et al., 2019).

Signals extracted from large blood vessels such as the internal carotid arteries and draining veins (superior sagittal sinus) have been shown to contribute systemic low-frequency oscillations to the fMRI signal that are vascular rather than neuronal in origin (Tong et al., 2019). Furthermore, the global signal is not spatially uniform and systemic low-frequency oscillations propagate through the brain with blood flow and arrive at different voxels at different times, rendering the standard regression approach temporally misaligned across space (Erdoğan et al., 2016). This tension between global signal as physiological confound and global signal as neural carrier has motivated data-driven criteria to determine when GSR is warranted, such as the Global Negative Index proposed by Chen et al. (2012), which estimates whether the global signal is dominated by neural or non-neural variance. However, such heuristics lack a principled algebraic foundation for GSR. Treating the global signal merely as a statistical nuisance variable or a proxy for arousal (Falahpour et al., 2018; Wong et al., 2013; Zhang & Northoff, 2022) obscures its underlying topological nature. While recent work has mapped the spatial topography of the global signal to specific systems like the default mode network (Bolt et al., 2022; Li, Bolt, et al., 2019), there remains a lack of rigorous theoretical frameworks that define exactly *what* topological component of the graph is being removed. The field lacks a precise geometric characterization of GSR that goes beyond its statistical effects on correlation matrices. This contentious discourse suggests the potential utility of a more general and geometrically-informed perspective rather than purely statistical characterization of GSR.

While GSR has traditionally referred to ‘Global Signal Regression’, the specific method of regressing out the global signal time course, in this paper we adopt the broader term ‘Global Signal Removal’ to encompass a family of spatial filtering operations that remove global signal variance through different mechanisms. We prove that the most common form of GSR, Regression-GSR, along with three other variants (Naive-GSR, PCA-GSR (Carbonell et al., 2011), and a new structure-informed variant we call SC-GSR), can all be expressed in the unified form 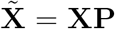 for spatial filter **P**. Each variant targets a different spatial mode: the uniform vector **1**_*N*_ (Naive), the first principal component **u**_1_ (PCA), the first structural harmonic ***ψ***_1_ (SC), or the functional degree vector **d** (Regression). This framework reveals key geometric distinctions between GSR variants, with implications for their effects on connectivity.

The paper is organized as follows. We begin by analyzing Regression-GSR, deriving its effect on the covariance matrix as a rank-1 update by the degree vector and recasting it as a spatial filter. We then show that the degree vector is dominated by the first eigenvector of the covariance matrix, linking Regression-GSR to principal component removal. Next, we extend the spatial filtering framework to three additional variants: Naive-GSR and PCA-GSR, which we recast in spatial filtering terms, and SC-GSR, a new variant that removes the first harmonic of the structural connectome. We demonstrate these theoretical results empirically using resting-state and task fMRI data from the Human Connectome Project (HCP), showing that the four variants form a spectrum from minimal distortion of the original pre-GSR connectivity to maximal reshaping of the connectome. Finally, we examine practical consequences: all variants induce numerical singularity, reshape resting-state network connectivity, and differentially interact with task-evoked signals. This work is intended as both a theoretical reference and a methodological guide, enabling investigators to select the spatial filter that best suppresses global signal while preserving task-relevant neural architecture.

## Regression-GSR

Let **X** ∈ ℝ^*T ×N*^ be the fMRI data matrix with *T* time points and *N* regions, where each regional time series is mean-centered: 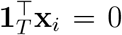 for *i* = 1, …, *N* . Throughout this paper, we use **1**_*K*_ to denote a *K* × 1 column vector of ones, and **I**_*K*_ to denote the *K* × *K* identity matrix.

The empirical covariance matrix, the most commonly used form of functional connectivity, is:

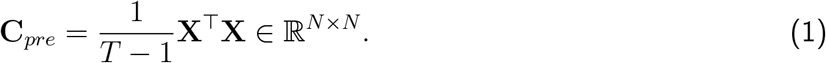

For standard Regression-GSR, the global signal **g** is defined as the spatial average time series:

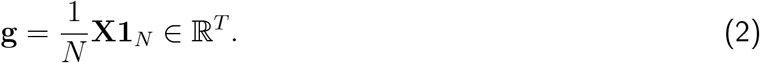

Regression-GSR projects the data onto the subspace orthogonal to **g** using the projection matrix:

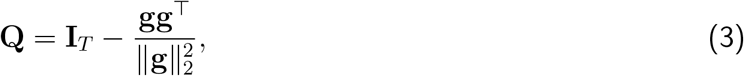

yielding the residualized data 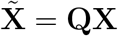. This operation projects the data onto the subspace orthogonal to the global signal, effectively removing any variance that correlates with **g**. The post Regression-GSR covariance matrix **C**_*post*_ is defined using the residualized data matrix 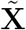 as

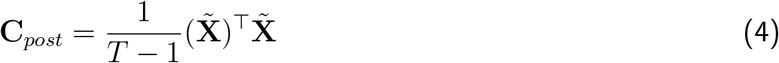

All propositions stated in this paper are proved in the Supplementary Material, with cross-references provided after each proposition.

### Regression-GSR projects out the covariance degree vector

To express the effect of Regression-GSR in terms of intrinsic network properties, we define the *degree vector* **d** = **C**_*pre*_**1**_*N*_, whose *i*-th entry *d*_*i*_ = ∑_*j*_ [**C**_*pre*_]_*ij*_ is the sum of all functional connections of region *i*.

#### Proposition 1

(Post Regression-GSR covariance as degree-based rank-1 update). *The effect of Regression-GSR on the covariance matrix is a rank-1 update determined entirely by the degree vector:*

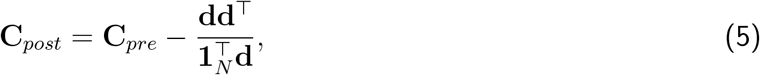

*where* 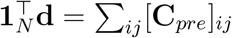 *is the sum of all entries in* **C**_*pre*_.

*Proof:* Proof of Proposition 1

#### Remark 1

(Interpretation of the rank-1 update). *Equation* (5) *reveals that GSR is not a uniform operation across brain regions. The element-wise change in covariance is* 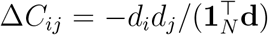, *which implies that connectivity reductions scale with the product of regional degrees. Hub regions (high d*_*i*_*) experience disproportionately large reductions, while peripheral regions remain relatively unaffected. This degree-dependent structure predicts that Regression-GSR will preferentially disrupt connectivity in hub-rich networks*.

#### Empirical support

Figure 1a illustrates an example of the covariance degree deflation underlying Regression-GSR (Equation 5), computed from resting-state fMRI data from Human Connectome Project (HCP) (Van Essen et al., 2013). **C**_*post*_ is obtained by subtracting a degree-weighted outer product from **C**_*pre*_. The covariance degree **d** exhibits spatial heterogeneity across resting-state networks, with higher values in sensory-motor systems (visual, somatomotor) and attention networks (dorsal attention, salience/ventral attention) compared to transmodal association cortex (default mode, frontoparietal).

**Figure 1.**
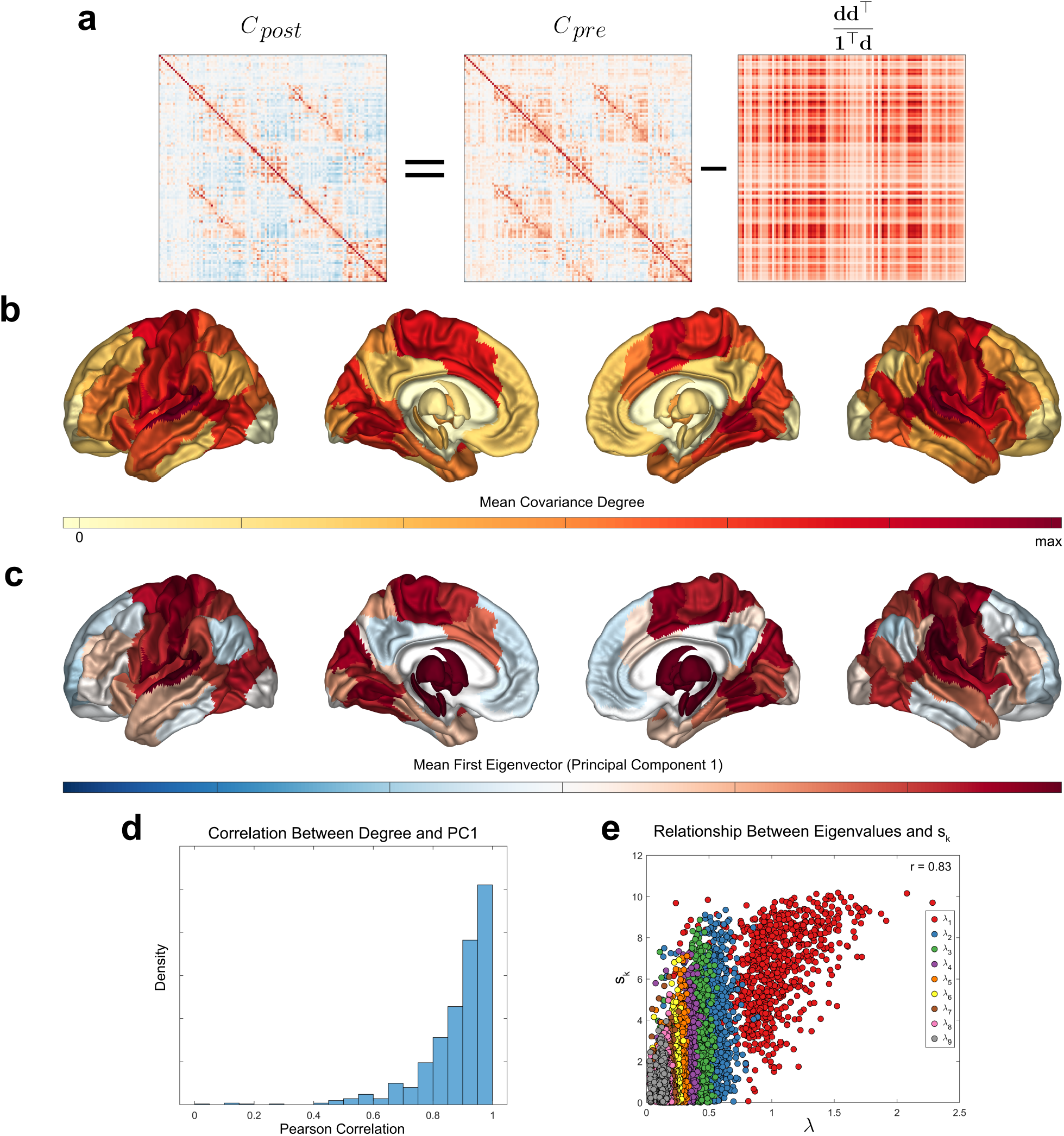
Regression-GSR overview: (a) An example of the covariance degree deflation underlying Regression-GSR (Equation 5). The covariance degree has varying influences across resting-state networks. (b) The group-level mean covariance degree in each brain region, and (c) the mean first eigenvector (i.e., principal component, “PC”) of the covariance matrix. (d) A histogram of the Pearson correlation between their covariance degree and the first PC across all subjects (mean ± std = 0.88±0.12). (e) The empirical relationship between *s*_*k*_ and *λ*_*k*_ across subjects (Pearson *r* = 0.83, *p <* 0.001).

### Eigenbasis representation of the covariance degree vector

In the previous section, we established that Regression-GSR performs a rank-1 update to the covariance matrix, subtracting a term proportional to **dd**^⊤^. To further understand the graph theoretic consequences of this operation, we now analyze the degree vector **d** in terms of the covariance eigenmodes and derive the effect of removing **d** on the eigenspectrum.

Let the eigendecomposition of the covariance matrix be:

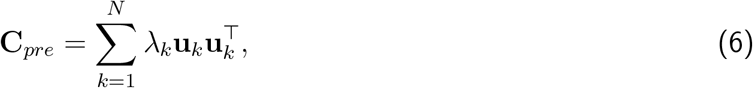

where *λ*_1_ ≥ *λ*_2_ ≥ · · · ≥ *λ*_*N*_ are the eigenvalues (representing the amount of variance captured by each eigenmode) and {**u**_*k*_} are orthonormal eigenvectors.

Substituting this decomposition into the definition **d** = **C**_*pre*_**1**_*N*_ yields the spectral expansion of the degree vector:

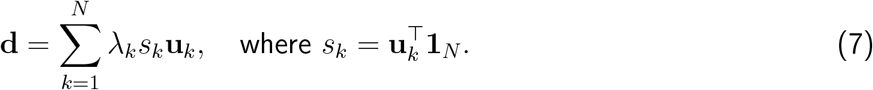

The scalar *s*_*k*_ quantifies the *global alignment* of the *k*-th eigenvector, measuring how uniformly its entries point in the same direction. Modes with all positive (or all negative) entries have large |*s*_*k*_|, while oscillatory modes with mixed signs have *s*_*k*_ ≈ 0 due to cancellation.

Equation (7) shows that **d** is a linear combination of all eigenvectors, with each contribution scaled by both variance (*λ*_*k*_) and global alignment (*s*_*k*_). However, in practice, a single term dominates this sum. For covariance matrices derived from fMRI data, the first eigenvector **u**_1_ captures the largest variance (*λ*_1_ ≫ *λ*_*k*_ for *k >* 1). Although covariance matrices can contain negative entries, fMRI covariance matrices are typically dominated by positive correlations, and in this regime the Perron-Frobenius theorem (Meyer, 2000) implies that the dominant eigenvector has all positive entries. Empirically, **u**_1_ is the only eigenvector with strictly positive entries in our data. Higher-order eigenvectors, being orthogonal to **u**_1_, must contain both positive and negative entries, driving their sums *s*_*k*_ toward zero.

Having established that **d** is dominated by the first eigenvector, we now characterize how the rank-1 update affects the full eigenspectrum. The post Regression-GSR covariance admits an eigendecomposition:

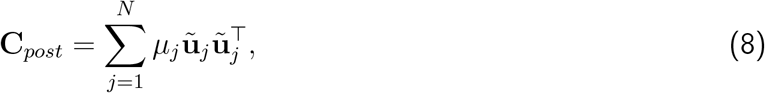

where *µ*_1_ ≥ *µ*_2_ ≥ · · · ≥ *µ*_*N*_ and {**ũ**_*j*_} are the post Regression-GSR eigenvalues and eigenvectors. Note that the indices *k* and *j* label eigenmodes in descending order of variance within each decomposition, but do not imply a correspondence between pre and post Regression-GSR eigenmodes. The rank-1 structure of GSR (Equation 5) renders the post Regression-GSR covariance singular, with a null space spanned by a direction determined by **d**. When **d** aligns closely with the dominant eigenvector **u**_1_, Regression-GSR approximately removes the largest eigenmode. The nonzero post Regression-GSR eigen-values satisfy interlacing inequalities with respect to the pre-GSR spectrum, though the correspondence between individual eigenmodes is generally not preserved.

In Supplementary Material, we provide a complete characterization of these spectral effects. Proposition S1 establishes that the eigenbasis is preserved when **d** aligns with a single eigenvector. Proposition S2 provides exact solutions for the post Regression-GSR eigenvalues via the secular equation. Proposition S3 proves eigenvalue interlacing, guaranteeing that all eigenvalues either decrease or remain unchanged. Proposition S4 shows that first-order eigenvalue shifts scale as 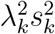, concentrating attenuation on the first eigenmode while leaving higher eigenmodes nearly unchanged.

#### Empirical support

We verify this spectral decomposition using resting-state fMRI data from the HCP young adult cohort (Figure 1). The spatial distribution of the covariance degree (Figure 1b) closely resembles the first eigenvector (Figure 1c), with both exhibiting elevated values in sensory-motor and attention networks. Quantitatively, the correlation corr(**d, u**_1_) averages 0.88 ± 0.12 across subjects (Figure 1d), confirming that **d** is highly correlated with **u**_1_. Notably, *λ*_*k*_ and |*s*_*k*_| are highly correlated across eigenmodes (Figure 1e; Pearson *r* = 0.83, *p <* 0.001), indicating that eigenvectors capturing the most variance are also the most globally aligned. This empirical property of fMRI covariance structure ensures that the spectral coefficients |*λ*_*k*_*s*_*k*_| concentrate heavily on the first mode. Thus, **d** is dominated by (but not identical to) the first eigenvector of the covariance matrix, with the strength of this approximation (corr(**d, u**_1_) ≈ 0.88) determining the extent to which Regression-GSR acts as clean first-mode removal versus a more complex reshaping of the full eigenspectrum.

### Regression-GSR can be expressed as a spatial filtering operation

Proposition 1 reveals what Regression-GSR removes (the degree vector), but not how it acts on the data. We now show that the standard regression procedure can be recast as a spatial filtering operation of the form 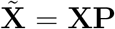, where **P** is a projection matrix acting on the spatial dimension. This reformulation connects the statistical regression coefficients directly to network topology.

#### Proposition 2

(Regression-GSR as graph spatial filtering). *Let* 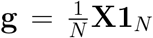 *be the global signal. In a standard regression framework, the residualized data is computed as* 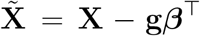, *where* ***β*** = **X**^⊤^**g***/*(**g**^⊤^**g**) *is the vector of regression coefficients. These statistical weights are strictly proportional to the covariance degree vector* **d** = **C**_*pre*_**1**_*N*_, *given by the exact relation:*

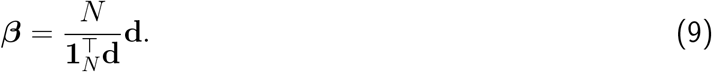

*By substituting this geometric relationship into the residual equation, we can eliminate the statistical coefficients and express Regression-GSR purely as a spatial filter acting on the covariance graph:*

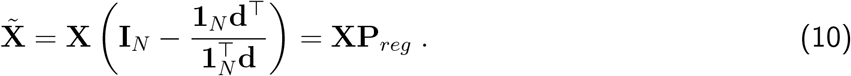

*The spatial filter* **P**_*reg*_ *has eigenvalues* {0, 1, …, 1}, *which eliminates the left and right eigenvectors with eigenvalue 0 corresponding to* **d**^⊤^ *and* **1**_*N*_ *(unnormalized), respectively. All other eigenvectors* **v**_*k*_ *satisfying* **d**^⊤^**v**_*k*_ = 0 *correspond to eigenvalue* 1.

*Proof:* Proof of Proposition 2

#### Remark 2

(Oblique projection onto degree-orthogonal subspace). *The spatial filter* **P**_*reg*_ *is an oblique projection (i*.*e*., *a non-orthogonal projection where the measurement and subtraction directions differ): it measures the projection onto* **d** *but subtracts along* **1**_*N*_ . *The regression coefficients* ***β*** ∝ **d** *weight each region by its connectivity degree, subtracting more from hub regions. The spatial filter satisfies* **d**^⊤^**P**_*reg*_ = **0**, *and at the covariance level this yields the rank-1 update* 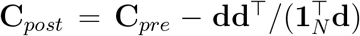 *established in Proposition 1*.

## Spatial filtering enables a family of GSR variants

Having established that Regression-GSR can be expressed as a spatial filter 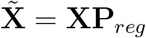 (Proposition 2), we now show that other GSR approaches fit within this same framework. Two existing methods, Naive-GSR, which removes the uniform vector **1**_*N*_, and PCA-GSR, which removes the first principal component **u**_1_, can both be written in the unified form 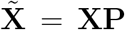. We also introduce a new variant, SC-GSR, which removes the first structural connectivity harmonic ***ψ***_1_. Unlike Regression-GSR, which is an oblique projection, these three variants are all orthogonal projections.

### Naive-GSR

The simplest GSR variant removes a uniform spatial average across all regions, treating each region identically regardless of its connectivity profile. The uniform vector **1**_*N*_ is the first eigenvector of the combinatorial graph Laplacian ℒ = **D** − **A** for any connected graph (i.e., a graph where a path exists between every pair of nodes) with adjacency matrix **A** and diagonal degree matrix **D**, giving Naive-GSR a natural interpretation as removal of the lowest-frequency graph mode.

#### Proposition 3

(Naive-GSR as graph spatial filtering). *Let* 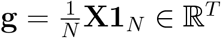 *be the global signal (spatial mean at each time point). Naive-GSR can be expressed as a spatial filter:*

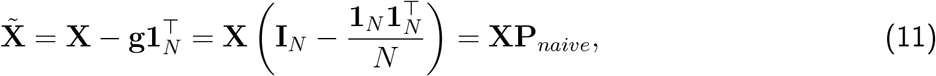

*where* **P**_*naive*_ *is an orthogonal projection onto the subspace orthogonal to* **1**_*N*_, *filtering out the homogeneous spatial contribution from all regions. The spatial filter* **P**_*naive*_ *has eigenvalues* {0, 1, …, 1} *with eigenvector* 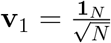 *corresponding to eigenvalue 0*.

*The post Naive-GSR covariance is:*

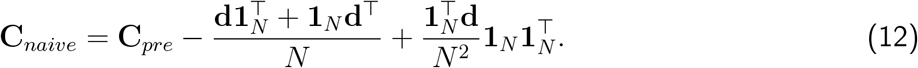

*Unlike Regression-GSR, which performs a rank-1 update, this is a rank-2 update in general. The two coincide when* **d** ∝ **1**_*N*_ . *Furthermore*, **C**_*naive*_**1**_*N*_ = **0**, *meaning the row sums (and by symmetry, column sums) of the post Naive-GSR covariance matrix are zero*.

*Proof:* Proof of Proposition 3

#### Remark 3

(Spatial high-pass filtering). *Naive-GSR is a spatial high-pass filter: the projection* **P**_*naive*_ *has eigenvalue 0 for* **1**_*N*_ *and eigenvalue 1 for all orthogonal modes, completely removing the spatially uniform component while preserving all other spatial patterns. This contrasts with Regression-GSR, which removes the degree vector* **d** *which is a data-dependent spatial mode shaped by network connectivity. At the covariance level, Naive-GSR performs a rank-2 update involving both* **d** *and* **1**_*N*_, *whereas Regression-GSR performs a rank-1 update along* **d** *alone*.

### PCA-GSR

An alternative approach directly targets the first principal component of the covariance matrix, removing the maximum variance mode (Carbonell et al., 2011; Leonardi et al., 2013).

#### Proposition 4

(PCA-GSR as graph spatial filtering). *Let* **C**_*pre*_ = **UΛU**^⊤^ *be the eigendecomposition of the pre-GSR covariance, with* **u**_1_ *the first eigenvector and λ*_1_ *the first eigenvalue. PCA-GSR can be expressed as a spatial filter:*

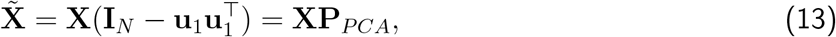

*where* **P**_*PCA*_ *is the spatial filter that eliminates* **u**_1_ *from the signal*.

*The post PCA-GSR covariance exactly removes the first eigenmode:*

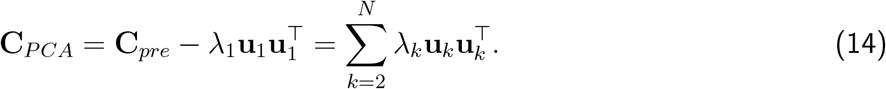

*Proof:* Proof of Proposition 4

#### Remark 4

(Exact eigenmode removal). *PCA-GSR removes the first eigenvector exactly, guaranteeing that no variance along* **u**_1_ *remains. This is also a rank-1 covariance update, but unlike Regression-GSR, it preserves the eigenbasis exactly: the post PCA-GSR eigenvectors are* {**u**_2_, …, **u**_*N*_ } *with eigenvalues* {*λ*_2_, …, *λ*_*N*_ }.

### SC-GSR

Recent work has investigated transforming functional brain signals into their representations of structural connectivity eigenmodes (or “harmonics”) (Abdelnour et al., 2018; Atasoy et al., 2016; Tewarie et al., 2020). Different structural harmonics appear to capture different processes and functional activity patterns (Preti & Van De Ville, 2019; Sipes et al., 2026). The connectome harmonics framework has an embedded notion of a global baseline mode: the first (or “trivial”) harmonic of the structural Laplacian. Building on this observation, we introduce SC-GSR, a new structure-informed variant that removes this first structural harmonic, targeting the anatomical degree distribution as a global signal to be removed from functional time series.

#### Proposition 5

(SC-GSR as graph spatial filtering). *Let* **G** *be the structural connectivity adjacency matrix (undirected, weighted)*, **D** *the degree matrix with D*_*ii*_ = ∑_*j*_ *G*_*ij*_, *and* ℒ_*SC*_ = **I** − **D**^−1*/*2^**GD**^−1*/*2^ *the symmetric normalized graph Laplacian. Let* ℒ_*SC*_ = **ΨΛ**_*SC*_**Ψ**^⊤^ *be its eigendecomposition. The first eigenvector is* ***ψ***_1_ = **D**^1*/*2^**1**_*N*_ */*∥**D**^1*/*2^**1**_*N*_ ∥ *with eigenvalue* 0 *(assuming the graph is connected)*.

*SC-GSR removes this first structural harmonic:*

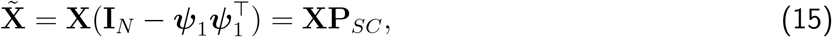

*where* **P**_*SC*_ *is the spatial filter that eliminates the structural connectome’s degree from the signal*.

*Proof:* Proof of Proposition 5

#### Remark 5

(First structural harmonic encodes degree). *The first eigenvector* ***ψ***_1_ ∝ **D**^1*/*2^**1**_*N*_ *directly encodes the structural degree distribution. Regions with high structural degree (anatomical hubs) have large entries in* ***ψ***_1_. *Thus, SC-GSR preferentially removes variance from hub regions*.

#### Empirical support

Having formalized the four GSR variants mathematically, we now compare them empirically using resting-state fMRI data from the HCP dataset (Figure 2). Figure 2a-b illustrates the progressive suppression of global co-fluctuations as one moves from the raw data to the aggressive PCA regression. To quantify this, we computed the Mean Squared Error (MSE) distance between approaches (Figure 2c) and extracted the dominant gradient of variation (Figure 2d). This analysis reveals a continuous “GSR Spectrum” bounded by No-GSR and PCA-GSR at opposite extremes. Crucially, the proposed SC-GSR removal clusters tightly with the naive and regression methods in the center of this spectrum.

**Figure 2.**
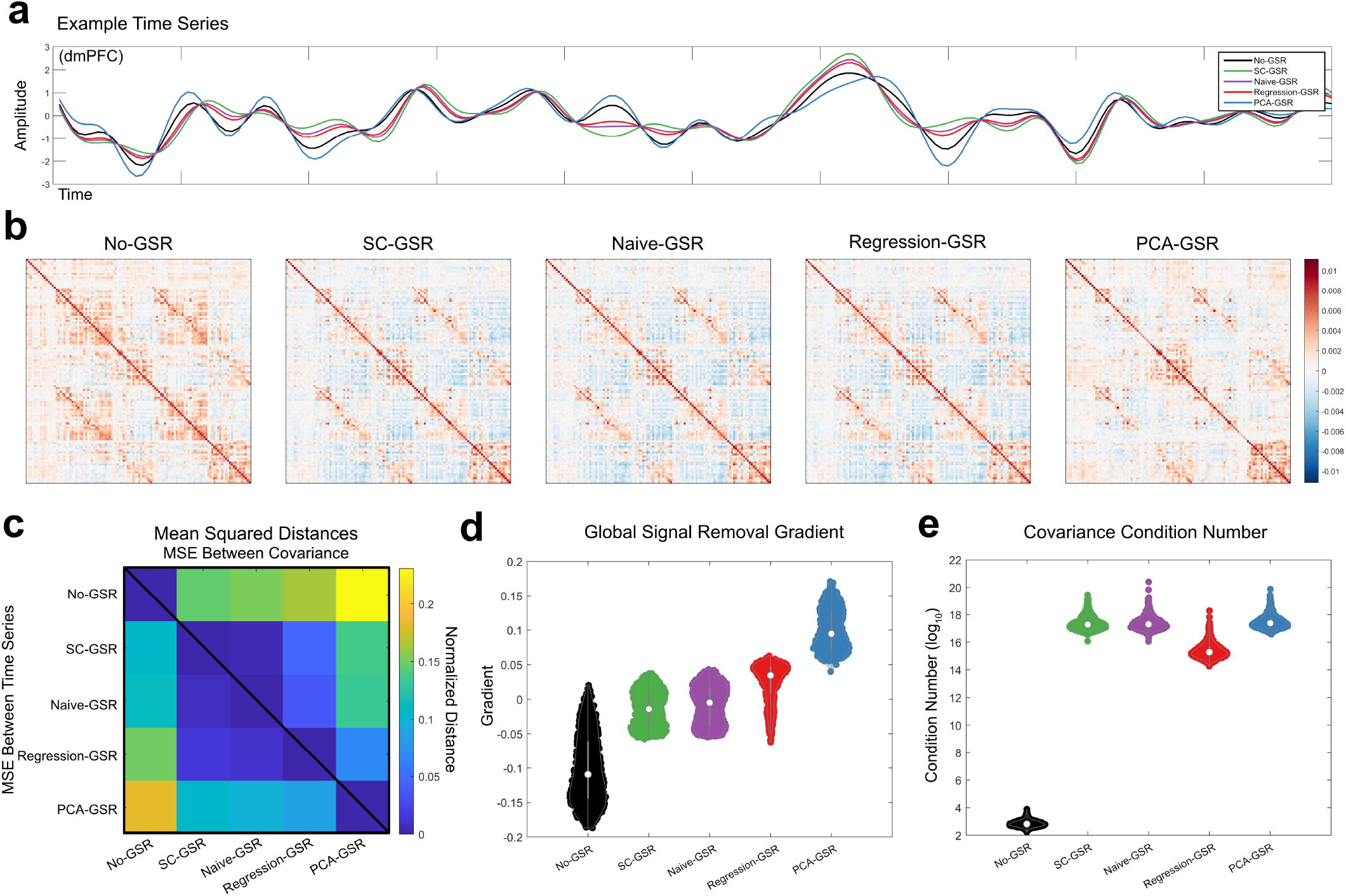
GSR variants: (a) A representative example fMRI time series showing the effects of each of four different GSR variants, along with the original time series (black). (b) A representative example of the different covariance matrices produced by different GSR variants. (c) A distance matrix with the lower-triangular indicating the group-averaged mean squared error (MSE) distance between time series of each variant and the upper-triangular indicating the group averaged MSE distance between covariance matrices for each variant. (d) Using the covariance MSE distance matrix, we performed classical multidimensional scaling to extract a 1-dimensional embedding (gradient) for the distances between each GSR variant for each subject, producing a group-level ordering of GSR variants. No-GSR and PCA-GSR live on opposite ends of this embedding, with the SC, Naive, and Regression-GSR variants living in-between them. (e) We calculated the condition number for each covariance matrix, plotting them on a log base-10 scale. GSR dramatically affects the conditioning of each covariance matrix.

## Practical consequences of GSR

The preceding sections formalized four distinct GSR variants, naive, regression, SC, and PCA, as a family of spatial filters, each targeting a specific topological feature of the connectome. We established that these variants are not arbitrary choices but form a deterministic geometric spectrum ranging from the uniform removal of the baseline (naive) to the targeted excision of the dominant eigenmode (PCA). We now examine how the choice of filter along this spectrum dictates the practical downstream effects on connectivity analysis. In this section, we demonstrate that moving along this “GSR Spectrum” systematically alters the numerical stability of covariance matrices, progressively reshapes the graph topology by inducing specific anticorrelations, and differentially modulates the separability of cognitive states. These results suggest that the “controversy” of GSR is better understood as a trade-off between maximizing signal retention (No-GSR) and enforcing topological orthogonality (PCA-GSR), with regression and SC-GSRs occupying a critical middle ground.

### GSR induces numerical singularity across all variants

Beyond its topological effects, GSR has immediate consequences for numerical stability. Many connectivity analyses, such as partial correlation and graphical LASSO, rely on calculating the precision matrix **C**^−1^. The reliability of this inversion is governed by the condition number *κ*(**C**) = *λ*_max_*/λ*_min_; a high *κ* indicates that the matrix is close to singular. Theoretically, because all GSR variants act as spatial projections (Propositions 2–5), they project the *N* -dimensional data onto a lower-dimensional subspace, mathematically forcing the smallest eigenvalue toward zero. Empirically, this effect is stark in the HCP resting-state data (Figure 2e): while unprocessed covariance matrices are well-conditioned (median *κ* ≈ 10^3^), all four GSR variants drive the condition number to ≈ 10^15^ or higher, approaching the limit of double-precision floating-point arithmetic.

This numerical fragility is particularly critical for *task-based fMRI*, where time series are often significantly shorter than in resting-state scans. When the number of time points (*T*) approaches the number of regions (*N*), the sample covariance matrix is inherently ill-conditioned even before preprocessing. In this regime, the pre-GSR matrices typically exhibit higher baseline condition numbers than their resting-state counterparts. Applying GSR which explicitly removes a dimension of variance compounds this issue, further destabilizing the eigenspectrum. Consequently, researchers analyzing short task runs with GSR cannot safely apply inverse-based metrics without aggressive regularization (e.g., shrinkage), as the resulting connectomes are effectively rank-deficient.

### GSR remodels connectivity in resting-state networks

Our geometric characterization reveals that GSR does not alter connectivity uniformly; rather, it remodels the network in a strictly degree-dependent manner. From Proposition 1, the change in covariance is given exactly by the rank-1 update 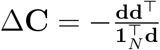. Element-wise, this means the connectivity between any two regions *i* and *j* is reduced by an amount proportional to the product of their weighted degrees:

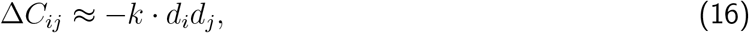

where *k* is a global scaling constant. This formulation leads to a specific topological prediction: “hub” regions (those with high degree **d**) will experience the most drastic reductions in covariance, effectively penalizing global integration to highlight local modularity.

#### Empirical support

We verified these predictions using resting-state fMRI data from the HCP young adult cohort, examining how GSR variants affect within- and between-network connectivity across eight canonical resting-state networks defined by the Schaefer parcellation (Figure 3). Without GSR, both within-network (dark colors) and between-network (light colors) covariances are predominantly positive across all networks, consistent with the global signal reflecting shared variance. All GSR variants significantly alter this structure (*p <* 0.001, Bonferroni corrected for all comparisons, paired t-test), but the spatial pattern of effects supports our degree-dependent framework.

**Figure 3.**
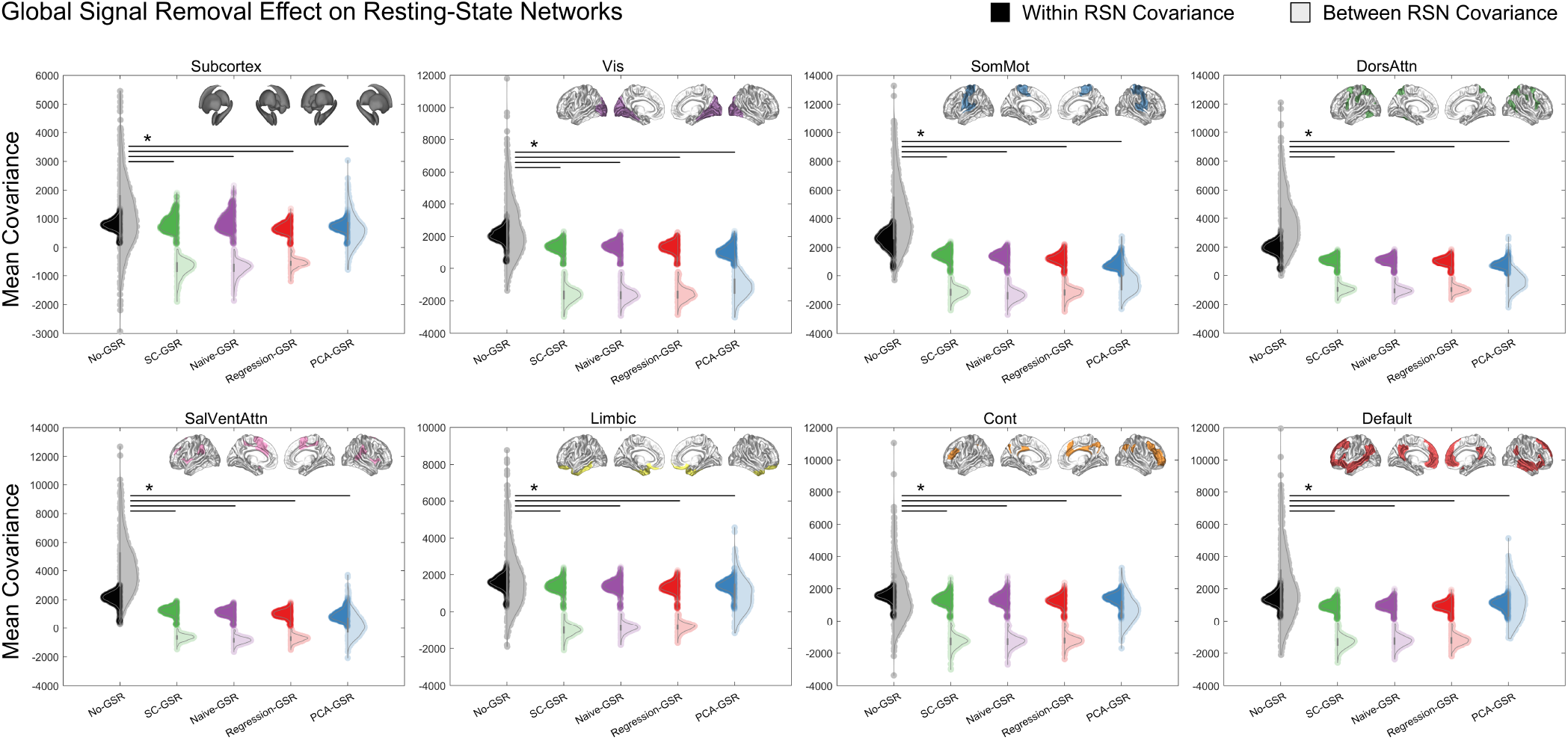
GSR remodels connectivity in resting-state networks. Within-network (dark colors) and between-network (light colors) mean covariance for eight canonical resting-state networks across GSR variants. Insets show the Schaefer parcellation used to define networks. All GSR variants significantly alter connectivity structure compared to No-GSR (*p <* 0.001, Bonferroni corrected for all comparisons, paired t-test). Between-network covariances shift strongly negative across all GSR variants, with SC, Naive, and Regression-GSR showing the most pronounced effects. Across GSR variants, within-network covariance reductions are largest in somatomotor and attention networks (dorsal attention, salience/ventral attention), and smallest in transmodal networks (default mode, control, limbic) and subcortical regions.

Between-network covariances shift strongly negative across all GSR variants, with SC, Naive, and Regression-GSR showing particularly pronounced negative shifts. This confirms that connections spanning different networks which involve regions with potentially different degree profiles experience systematic deflation that often exceeds the original positive correlation, inducing anticorrelations. PCA-GSR shows similar but slightly attenuated effects, reflecting the difference between removing **d** versus **u**_1_. Within-network covariances show heterogeneous effects. Somatomotor, salience/ventral attention, and dorsal attention networks show the largest absolute reductions in within-network covariance, while transmodal networks (default mode, control, limbic) and subcortical regions show smaller reductions. Visual network shows an intermediate pattern. For Regression-GSR, this aligns with the degree-weighted deflation in Proposition 1: connectivity reductions scale with the product of regional degrees, so networks with higher within- or between-network degree sums experience larger reductions.

### GSR in task fMRI

Our spectral interpretation rests on a key assumption: the degree vector is dominated by the first eigenvector, **d** ≈ *λ*_1_*s*_1_**u**_1_. When this holds, Regression-GSR approximates removal of the first principal component which is the global baseline mode. However, task-evoked activity can violate this assumption in two ways. First, cognitive tasks engage specific functional networks, potentially increasing their variance relative to the global mode. If a task network’s variance approaches or exceeds *λ*_1_, the eigenvector ordering may shift, and GSR would target the task-dominant network rather than the global baseline. Second, the degree vector may overlap substantially with task-activated regions. Let ***b*** be a map of General Linear Model (GLM) regression coefficients identifying brain regions responding to a task design. For each GSR variant, we define

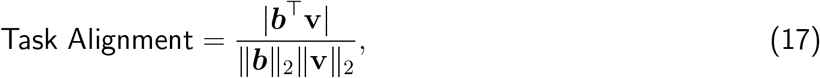

which measures the cosine similarity between the task activation pattern and the GSR target vector **v** (i.e., **d** for Regression-GSR, **1**_*N*_ for Naive-GSR, **u**_1_ for PCA-GSR, or ***ψ***_1_ for SC-GSR). High Task Alignment indicates that the GSR variant will preferentially remove variance from task-relevant regions, potentially eliminating task-related signal.

#### Empirical support

We evaluated these concerns using task fMRI data from seven HCP tasks (emotion, motor, gambling, language, social, relational, and working memory) alongside resting-state data. First, we quantified the “risk of signal loss” by computing the Task Alignment (cosine similarity) between each method’s filter vector and the task-activation maps (**b**) derived from a standard GLM analysis (Figure 4). The results reveal a stark dichotomy in how GSR variants interact with task structure. Standard Regression and PCA-GSR exhibited significantly higher alignment with task activation vectors, particularly in cognitively demanding tasks such as working memory, gambling, relational, and social (*p <* 0.05, FDR corrected, paired t-test). This confirms that for data-driven subtractive methods, the “global noise” being removed is often collinear with the “global signal” of the task itself. In contrast, SC and Naive-GSR showed significantly lower alignment across all tasks (*p <* 0.05, FDR corrected, paired t-test). By anchoring the removal to stable anatomical or uniform modes rather than transient functional covariance, these methods are less likely to “chase” and excise task-evoked variance.

**Figure 4.**
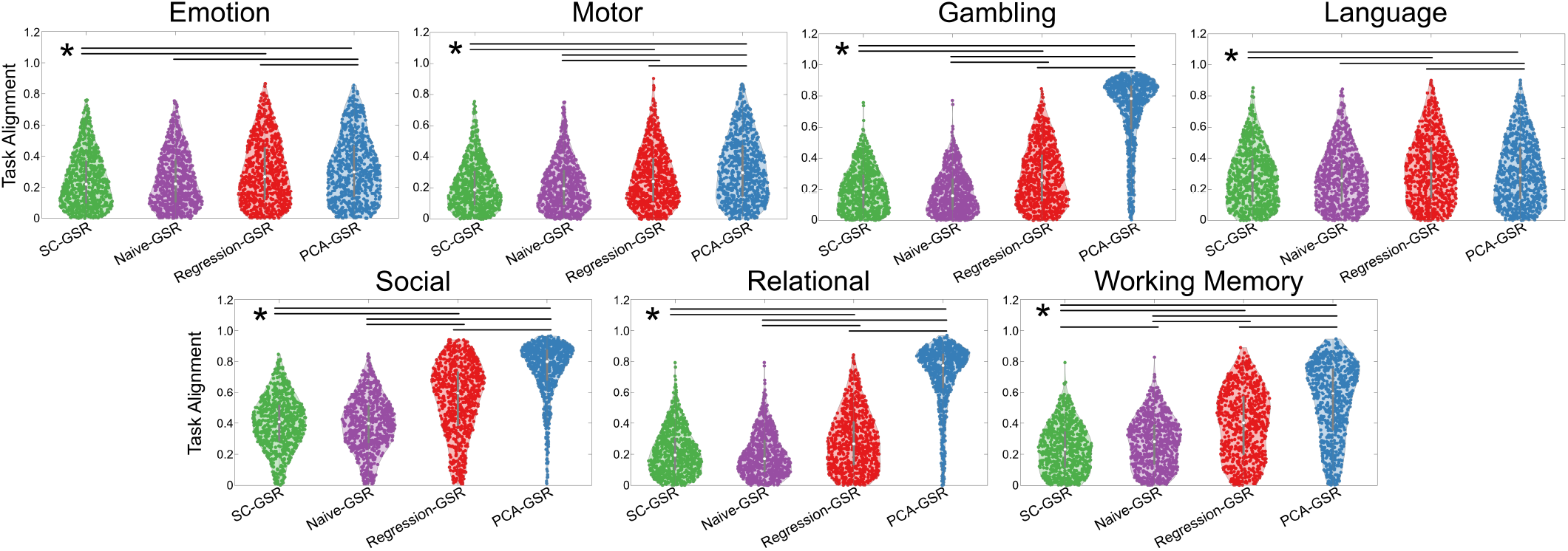
Task Alignment across GSR variants. Each subject’s task-activation vector **b** was calculated using a standard General Linear Model (GLM) approach. Task Alignment (Equation 17) measures the cosine similarity between **b** and each GSR variant’s target vector. SC-GSR and Naive-GSR show significantly lower Task Alignment across all tasks, while Regression-GSR and PCA-GSR show higher Task Alignment, particularly for cognitively demanding tasks (working memory, gambling, relational, social). Horizontal lines indicate significant pairwise differences (*p <* 0.05, FDR corrected, paired t-test).

We then assessed the downstream impact on task-state separability using t-Distributed Stochastic Neighbor Embedding (t-SNE, perplexity = 30) on two connectivity measures: covariance degree and positive FC degree (Figure 5). The results reveal distinct effects depending on the measure. For covariance degree (Figure 5a), Naive and Regression-GSR produce **d** = **0** by construction, yielding uninformative embeddings. In contrast, SC-GSR preserves non-trivial covariance degree structure and achieves the highest silhouette scores (using squared Euclidean distance) among all variants, though task separation remains imperfect. For positive FC degree (Figure 5b), SC, Naive, and Regression-GSR all enhance task clustering compared to No-GSR and PCA-GSR. These results suggest that SC-GSR best preserves task-specific covariance structure, while multiple GSR variants (SC, Naive, Regression) enhance task separability in positive FC degree patterns.

**Figure 5.**
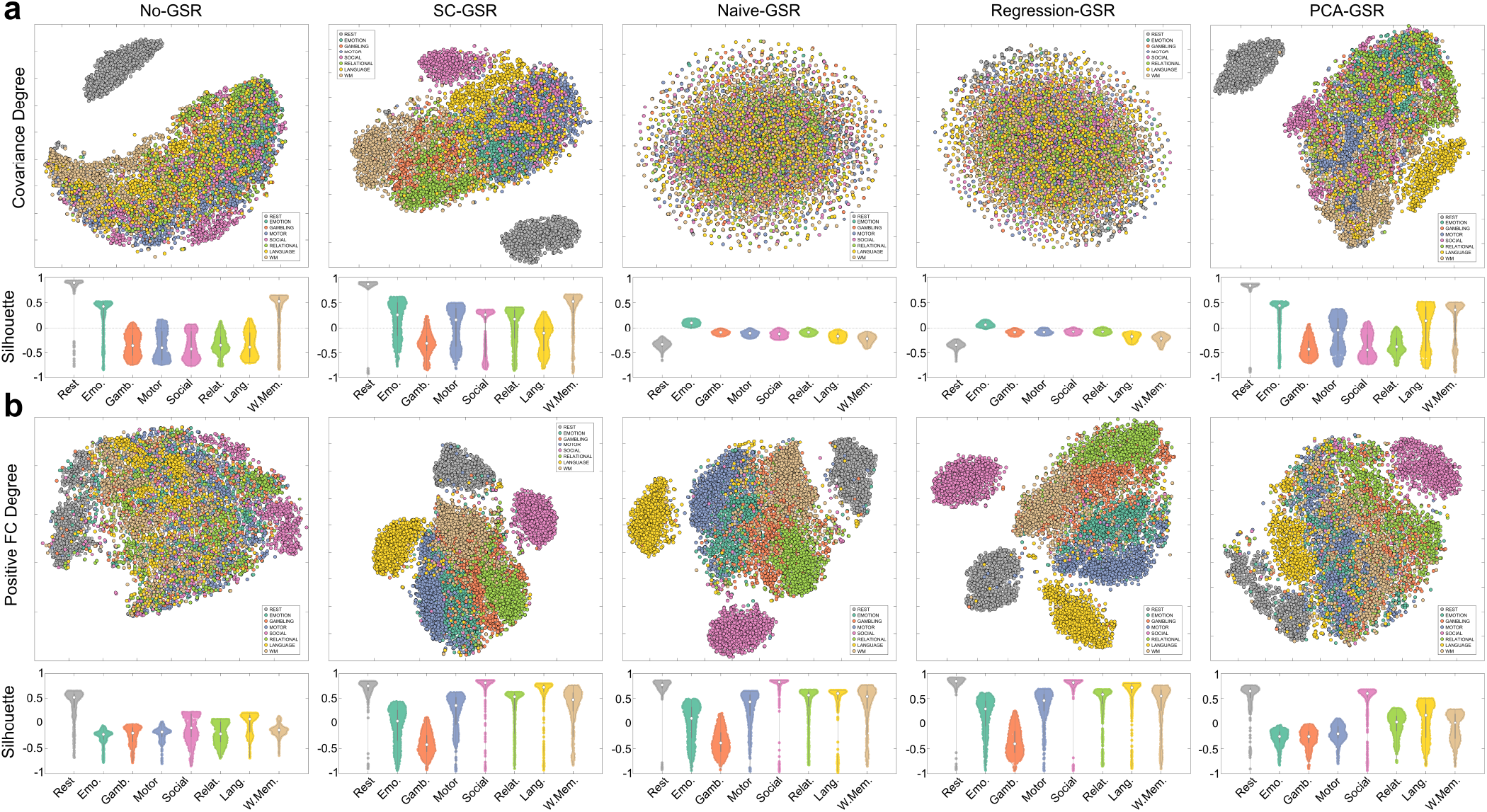
GSR variants differentially affect task-state separability. t-SNE embeddings of (a) covariance degree and (b) positive FC degree across GSR variants, color-coded by task. Squared Euclidean distance silhouette scores (violin plots) quantify task clustering quality: higher values indicate better-separated, more tightly clustered tasks. For covariance degree, SC-GSR produces the best task separation, while Naive and Regression-GSR produce zero covariance degree by construction (**d** = **0**), yielding uninformative random embeddings shown for completeness. For positive FC degree, SC, Naive, and Regression-GSR all enhance task clustering compared to No-GSR and PCA-GSR. Results indicate that removing structural baseline (SC harmonics) optimally preserves task-specific covariance signatures, while removing global mean (Regression, Naive) better preserves task structure in positive FC degree patterns.

## Datasets and preprocessing

We validated our theoretical predictions using 770 subjects resting-state fMRI and diffusion MRI data from the Human Connectome Project (HCP) S1200 release (Van Essen et al., 2013) ^1^. We included all HCP subjects that had both diffusion and functional MRI, and excluded subjects with quality control issues. Unprocessed HCP data were downloaded and organized into BIDS format. Anatomical scans were collected with an MPRAGE T1-weighted sequence. Functional MRI (fMRI) data included four 15-minute rest sessions and two task sessions for seven tasks (emotion, motor, gambling, language, social, relational, and working memory), all with a repetition time (TR) of 720 ms. Additionally, we used the pre-processed diffusion tensor imaging (DTI) data made available by HCP, which includes merging high fidelity directions across multiple multi-shell DTI acquisitions (b = 1000, 2000, & 3000) and correcting for phase-encoding polarity distortion (Glasser et al., 2013). We applied a state-of-the-art image processing pipeline, micapipe (Cruces et al., 2022), which processes the anatomical, functional, and diffusion MRI data in a coherent framework to produce subject-specific structural connectivity and functional regional time series in the Schaefer atlas with 100 cortical regions (Schaefer et al., 2018), each with connectivity to an additional 14 subcortical regions (Patenaude et al., 2011).

The structural connectivity was computed using MRtrix3 and probabilistic tractography, generating 10 million streamlines throughout the gray matter-white matter interface (maximum tract length = 400, minimum length = 10, cutoff = 0.06, step = 0.5, angle curvature = 22.5^*◦*^) using the iFOD2 algorithm and 3-tissue anatomically constrained tractography (Cruces et al., 2022). Tractograms were filtered using SIFT2 (Smith et al., 2015), reweighting streamlines by cross-sectional multipliers to provide connection densities that are biologically valid measures of fiber connectivity.

The fMRI underwent image re-orientation, motion and distortion correction, and nuisance signal regression (white matter, cerebrospinal fluid (CSF), and frame-wise displacement spikes). Volumetric time series were mapped to the native FreeSurfer space using boundary-based registration (Greve & Fischl, 2009). Vertex-level time series were averaged into parcels defined by the Schaefer atlas with 100 cortical regions (Schaefer et al., 2018). Parcel-level time series were low-pass filtered to be less than 0.1Hz, then underwent GSR preprocessing (with variants as described in the main text).

## Discussion

Global Signal Removal (GSR) has historically been debated through the lens of signal origin: whether the global signal represents physiological artifact or meaningful neural activity. Our work reframes this controversy by providing a precise linear algebraic formalization: all GSR variants can be expressed as spatial filters of the form 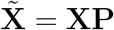, each targeting a different spatial mode. Naive-GSR, PCA-GSR, and SC-GSR are orthogonal projections removing the uniform vector, the first functional eigenvector, and the first structural harmonic, respectively.

Regression-GSR is an oblique projection that measures along the covariance degree vector but subtracts along the uniform vector, producing a rank-1 covariance update. Because the degree vector is dominated by the first eigenvector of functional connectivity, Regression-GSR approximately removes this dominant mode. The covariance change 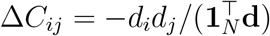 shows that Regression-GSR penalizes connectivity in a strictly degree-dependent manner: connections between high-degree regions experience the largest absolute reductions. The other three variants act similarly as spatial filters removing a global baseline mode, but with different spatial targets: the uniform vector (Naive), the first functional eigenvector (PCA), or the first structural harmonic (SC).

This topological remodeling introduces critical trade-offs for statistical stability and network interpretation. We demonstrate that all GSR variants drive the covariance condition number toward numerical singularity, effectively rendering the post-GSR connectome rank-deficient. This fragility necessitates the use of regularization (e.g., shrinkage) for any inverse-based analyses, particularly in short task-based scans where the empirical covariance matrix is already ill-conditioned. Furthermore, the choice of filter variant dictates the resulting network structure: empirically, all variants shift between-network correlations negative, with Regression, Naive, and SC-GSR showing the most pronounced effects and PCA-GSR showing slightly attenuated shifts. Within-network effects vary by network type: somatomotor and attention networks show the largest reductions, whereas transmodal networks (default mode, control, limbic) and subcortical regions show smaller reductions. Researchers must therefore choose their variant based on their specific hypothesis.

Finally, the correct application of GSR to task fMRI requires a departure from uniform “on/off” pre-processing pipelines in favor of a spatially targeted approach. Our analysis reveals that for cognitively demanding tasks (e.g., working memory, gambling, relational, and social), the target spatial vectors of data-driven methods (Regression-GSR and PCA-GSR) significantly align with task-evoked activation maps (*p <* 0.05, FDR corrected, paired t-test). This alignment indicates a risk of removing task-relevant variance. In contrast, methods anchored to anatomical modes, such as SC-GSR, show lower alignment with task activations. The consequences for task-state separability depend on the connectivity measure: for covariance degree, Naive and Regression-GSR produce uninformative embeddings (**d** = **0** by construction), while SC-GSR preserves task-specific structure. For positive FC degree, SC, Naive, and Regression-GSR all enhance task clustering compared to No-GSR and PCA-GSR. We therefore propose that researchers explicitly validate their chosen GSR variant by inspecting the projection of its specific target spatial vector (e.g., the degree vector for Regression-GSR, or the first eigenvector for PCA-GSR) onto the task-evoked activation maps (e.g., GLM weights). When the GSR target vector and the task activations are highly collinear, that specific GSR method should be avoided or replaced with anatomically constrained alternatives. One limitation of this work is that we did not systematically evaluate interactions between GSR and other preprocessing steps such as motion correction or temporal filtering, which may modulate some of the effects described here.

## Conclusion

This work reframes Global Signal Removal (GSR) from a statistical correction into a deterministic spatial filtering operation. By mapping GSR onto the brain’s graph spectral embedding, we demonstrate that it acts as a high-pass graph filter, suppressing the baseline of global integration coupled to the brain’s structural hubs. We introduce SC-GSR, a new structure-informed variant that removes the first harmonic of the structural connectome, providing an anatomically grounded alternative to data-driven methods. While all GSR variants, including Naive, Regression, PCA, and SC-GSR, render covariance matrices numerically singular, they differ critically in their selectivity. Data-driven methods like standard regression and PCA aggressively excise variance that often overlaps with cognitive task activation, risking signal loss, whereas anatomically constrained variants like SC-GSR better preserve task-specific connectivity signatures. Consequently, GSR should be treated as a hypothesis-driven choice to penalize global integration in favor of local modularity, requiring researchers to select the spatial filter that best aligns with the specific neural phenomenon they seek to isolate.

## Data and code availability

All data used in this study are publicly available from the Human Connectome Project. All source code for the empirical supports will be made publicly available on GitHub upon publication.

## Author contributions

F.A. and B.S.S. contributed equally to this work. F.A. and B.S.S. developed the theoretical framework and performed the analyses. S.S.N. and A.R. supervised the project. All authors contributed to writing and reviewing the manuscript.

## Acknowledgments

This work is supported by NIH grants R01AG072753, R21AG087921, RF1AG087302, RF1NS100440.

## Supplementary Material

### Regression-GSR selectively reshapes the covariance eigenstructure

We analyze how the rank-1 update in Proposition 1 affects the eigenstructure of the covariance matrix. We first show that Regression-GSR can induce eigenvector rotations, but these rotations are negligible when the degree vector is dominated by the first eigenvector, a condition satisfied in resting-state data where corr(**d, u**_1_) ≈ 0.88.

#### Proposition S1

(Effect of GSR on covariance eigenvectors). *Let* **C**_*pre*_ = **UΛU**^⊤^ *be the eigendecomposition, where* **U** = [**u**_1_| · · · |**u**_*N*_ ] *contains orthonormal eigenvectors and* **Λ** = diag(*λ*_1_, …, *λ*_*N*_) *contains eigenvalues. Define* **s** = **U**^⊤^**1**_*N*_, *the vector of global alignments, with* 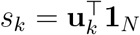. *Then:*

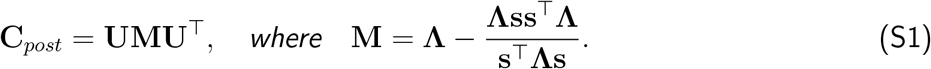

*The matrix* **M** *represents* **C**_*post*_ *in the pre-GSR eigenbasis, with elements:*

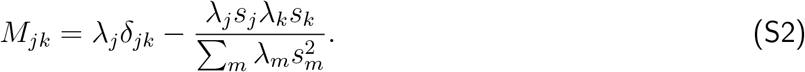

*The matrix* **M** *is diagonal if and only if* **d** ∥ **u**_*k*_ *for some k. Consequently:*

1. *In general*, **M** *is not diagonal, meaning GSR alters both the eigenvalues and eigenvectors of the co-variance matrix. The post Regression-GSR eigenvectors* {**ũ**_*j*_} *differ from the pre-GSR eigenvectors* {**u**_*k*_}.
2. *If* **d** ∥ **u**_1_, *then* **M** = diag(0, *λ*_2_, …, *λ*_*N*_). *Since* **M** *is diagonal, the post Regression-GSR eigenvectors* 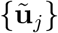 *coincide with the pre-GSR eigenvectors* {**u**_*k*_}. *The post Regression-GSR eigenvalues are* {0, *λ*_2_, …, *λ*_*N*_ }: *the eigenvalue associated with* **u**_1_ *becomes zero, while the eigenvalues associated with* **u**_*k*_ *for k* ≥ 2 *remain unchanged at λ*_*k*_.

*Proof:* Proof of Proposition S1

Thus Regression-GSR approximately preserves the eigenbasis while selectively modifying eigenvalues. We focus here on characterizing these eigenvalue changes, as they determine the practical consequences for connectivity analysis.

We denote the pre-GSR eigenvalues as *λ*_1_ ≥ *λ*_2_ ≥ · · · ≥ *λ*_*N*_ with corresponding eigenvectors **u**_1_, …, **u**_*N*_ . The post Regression-GSR eigenvalues, ordered by magnitude, are denoted *µ*_1_ ≥ *µ*_2_ ≥ · · · ≥ *µ*_*N*_, with corresponding eigenvectors **ũ**_1_, …, **ũ**_*N*_ . Note that *µ*_*j*_ is the *j*-th largest post Regression-GSR eigenvalue, which may not correspond to pre-GSR eigenvector **u**_*j*_ if the spectrum reorders.

Since Proposition 1 expresses **C**_*post*_ as a rank-1 update to **C**_*pre*_, the new eigenvalues *µ*_1_, …, *µ*_*N*_ can be characterized exactly using classical results on rank-1 perturbations (Golub & Van Loan, 2013).

#### Proposition S2

(Secular equation for post Regression-GSR eigenvalues). *The eigenvalues µ*_1_, …, *µ*_*N*_ *of* **C**_*post*_ *are the roots of the secular equation:*

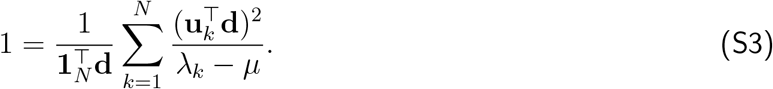

*Proof:* Proof of Proposition S2

#### Proposition S3

(Eigenvalue interlacing). *The new eigenvalues interlace with the original eigenvalues:*

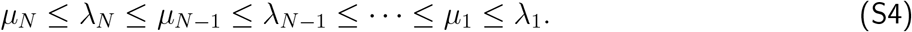

*That is, exactly one new eigenvalue lies in each interval* (−∞, *λ*_*N*_ ], [*λ*_*N*_, *λ*_*N*−1_], …, [*λ*_2_, *λ*_1_].

*Proof:* Proof of Proposition S3

#### Proposition S4

(First-order eigenvalue perturbation). *Under GSR, the eigenvalue associated with eigen-vector* **u**_*k*_ *admits the perturbation expansion* 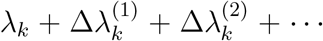, *where the first-order term is exactly:*

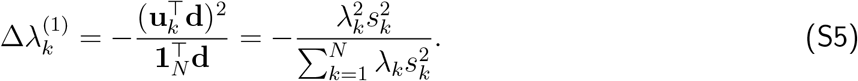

*When eigenvalue gaps are large relative to the perturbation magnitude, higher-order terms are negligible*.

*Proof:* Proof of Proposition S4

#### Remark S1

(Interpretation of eigenvalue effects). *Propositions S2, S3, and S4 provide complementary characterizations of eigenvalue changes. Proposition S2 gives the secular equation for the exact post Regression-GSR eigenvalues. Only modes with nonzero projection onto* **d** *contribute to this equation; eigenvectors orthogonal to* **d** *are completely unaffected by GSR. Proposition S3 establishes interlacing inequalities guaranteeing that the j-th largest post Regression-GSR eigenvalue is bounded by consecutive pre-GSR eigenvalues, with exactly one post Regression-GSR eigenvalue in each interval between consecutive pre-GSR eigenvalues. Proposition S4 shows that first-order shifts scale as* 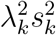, *concentrating attenuation on the first eigenvalue while leaving higher eigenvalues nearly unchanged. In the limit* **d** ∥ **u**_1_, *the eigenvalue associated with* **u**_1_ *vanishes entirely (becoming the smallest eigenvalue), while eigenvalues associated with* **u**_*k*_ *for k >* 1 *remain at λ*_*k*_.

### Proof of Proposition S1

We express the post Regression-GSR covariance matrix in the eigenbasis of **C**_*pre*_ and determine when the resulting matrix is diagonal.

We can first express vector **d** in the covariance eigenbasis as:

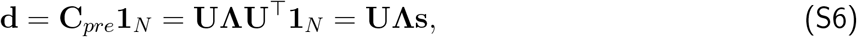

where **s** = **U**^⊤^**1**_*N*_ is the vector of projections of **1**_*N*_ onto each eigenvector, i.e., 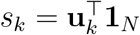. The denominator 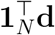 can be written as:

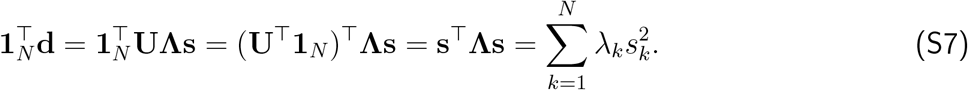

The numerator **dd**^⊤^ can also be written as:

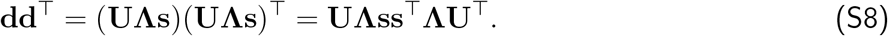

Substituting into the GSR formula from Proposition 1, 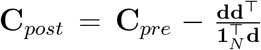, we can write the post Regression-GSR covariance as:

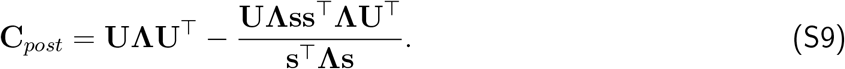

Factoring out **U** and **U**^⊤^:

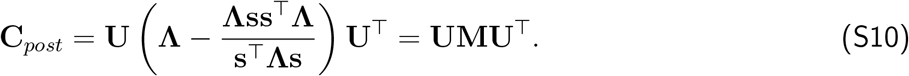

The matrix **M** = **U**^⊤^**C**_*post*_**U** represents **C**_*post*_ in the original eigenbasis. Its (*j, k*) element is:

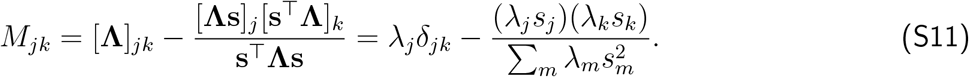

The off-diagonal elements are:

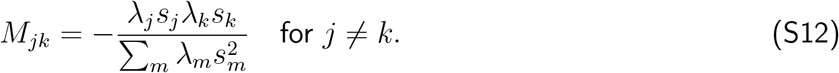

These vanish for all *j*≠ *k* if and only if at most one element of the vector (*λ*_1_*s*_1_, *λ*_2_*s*_2_, …, *λ*_*N*_ *s*_*N*_) is nonzero. Since *λ*_*k*_ *>* 0 for all *k* (assuming **C**_*pre*_ is positive definite), this requires at most one *s*_*k*_≠ 0. Since **d** = **UΛs** = ∑_*k*_ *λ*_*k*_*s*_*k*_**u**_*k*_, having exactly one nonzero *s*_*k*_ is equivalent to **d** ∥ **u**_*k*_ for some *k*.

For the special case of **d** ∥ **u**_1_, we have *s*_*k*_ = 0 for all *k* ≥ 2, then **s** = (*s*_1_, 0, …, 0)^⊤^. The denominator becomes:

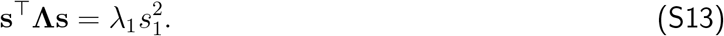

The matrix **Λss**^⊤^**Λ** has only one nonzero element:

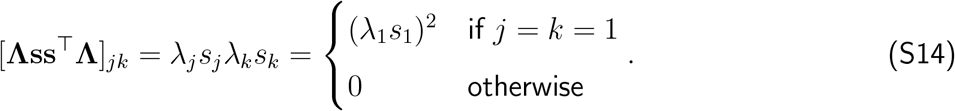

Therefore:

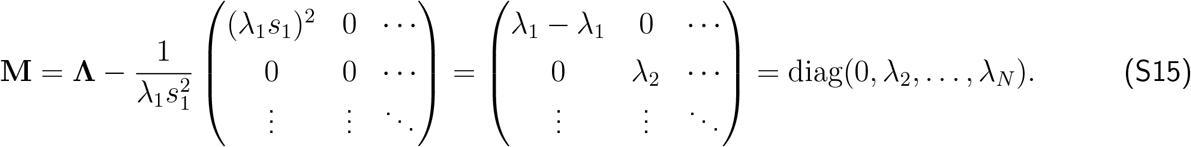

Since **M** is diagonal, the post Regression-GSR eigenvectors {**ũ**_*j*_} coincide with the pre-GSR eigenvectors {**u**_*k*_}. The post Regression-GSR eigenvalues are {0, *λ*_2_, …, *λ*_*N*_ }: the eigenvalue associated with **u**_1_ is zero, while eigenvalues associated with **u**_*k*_ for *k* ≥ 2 remain at *λ*_*k*_.

### Proof of Proposition S2

The eigenvalues of **C**_*post*_ satisfy det(**C**_*post*_ − *µ***I**) = 0. Substituting **C**_*post*_ = **C**_*pre*_ − *c***dd**^⊤^:

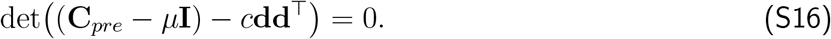

By the matrix determinant lemma, for any invertible matrix **A** and vector **v**:

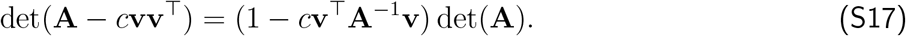

Applying this with **A** = **C**_*pre*_ − *µ***I** and **v** = **d**:

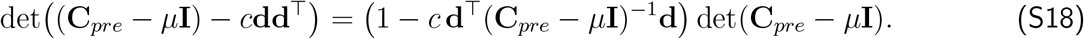

For this product to equal zero, either det(**C**_*pre*_ − *µ***I**) = 0, which gives the original eigenvalues *λ*_*k*_, or the first factor vanishes:

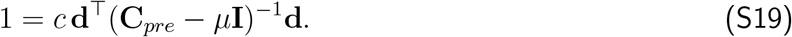

To evaluate the quadratic form, we use the eigendecomposition 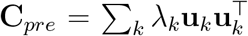. Since the eigen-vectors form an orthonormal basis, 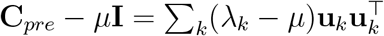, and thus:

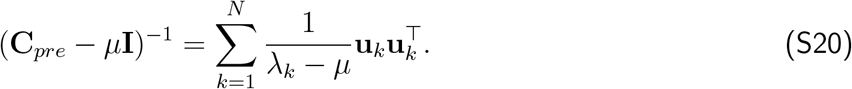

The quadratic form becomes:

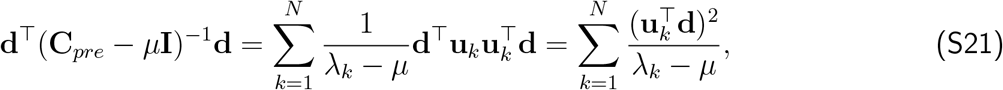

where we used 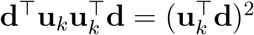. Substituting 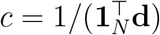 yields equation (S3).

### Proof of Proposition S3

Define 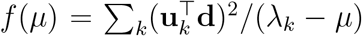. The secular equation (S3) can be written as 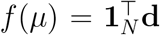. The function *f* has three key properties: (i) vertical asymptotes at each *λ*_*k*_ where *f* → +∞ from the left and *f* → −∞ from the right; (ii) *f* (*µ*) → 0^+^ as *µ* → −∞; and (iii) *f* is strictly increasing between asymptotes since 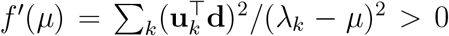. Since 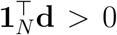, the horizontal line 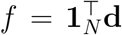 intersects *f* (*µ*) exactly once in each interval between consecutive asymptotes, yielding the interlacing pattern.

Alternatively, since *c***dd**^⊤^ ⪰ 0 (positive semi-definite), Weyl’s inequality implies that the *j*-th largest eigenvalue of **C**_*post*_ is bounded above by the *j*-th largest eigenvalue of **C**_*pre*_, i.e., *µ*_*j*_ ≤ *λ*_*j*_ for all *j*.

### Proof of Proposition S4

For a symmetric matrix **A** with non-degenerate eigenpair (*λ*_*k*_, **u**_*k*_), standard perturbation theory states that under a symmetric perturbation **A** → **A** + **E**, the first-order eigenvalue shift is 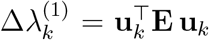.

Since 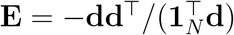 is symmetric:

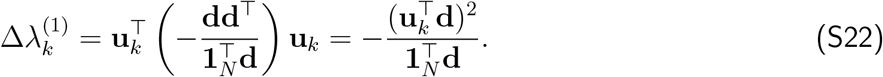

Substituting 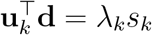 from equation (7) and 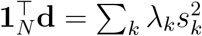 yields the final form.

### Proof of Proposition 1

Since **Q** is an orthogonal projection (symmetric and idempotent: **Q**^⊤^ = **Q** and **Q**^2^ = **Q**), the post Regression-GSR covariance is:

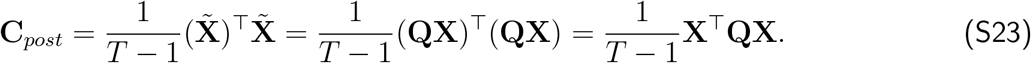

Substituting the residual-maker matrix 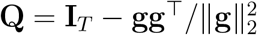 and expanding:

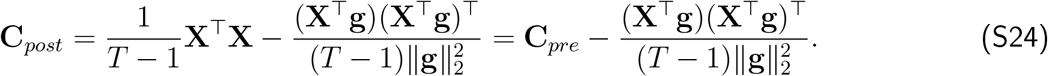

We now express the quantities **X**^⊤^**g** and 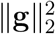 in terms of **d**. From equations (1) and (2):

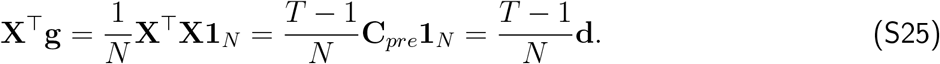

Similarly:

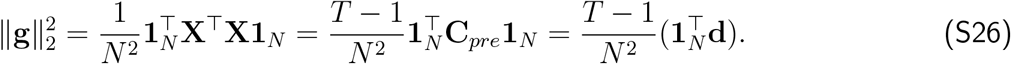

Substituting into equation (S24):

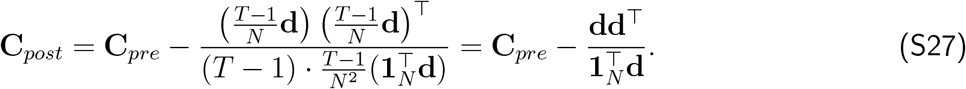

### Proof of Proposition 2

We first show that the regression coefficients ***β*** are proportional to the degree vector. Substituting 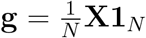 into ***β*** = **X**^⊤^**g***/*(**g**^⊤^**g**):

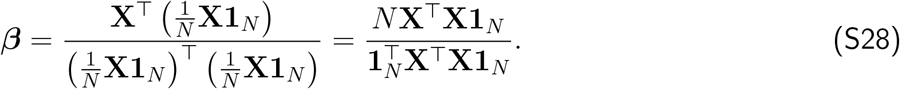

Since 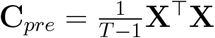 and **d** = **C**_*pre*_**1**_*N*_, the factor (*T* − 1) cancels out:

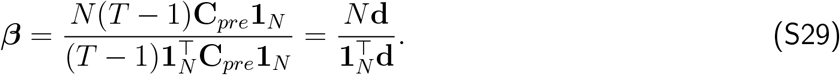

We next derive the spatial filter form by substituting both 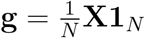 and 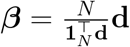 into the temporal residual equation:

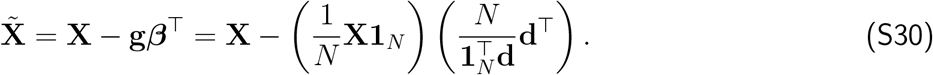

The *N* terms cancel perfectly, yielding the pure spatial filter:

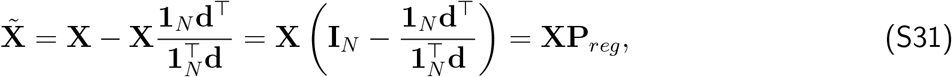

Where 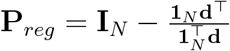.

We now examine the eigenstructure of **P**_*reg*_. We note that the scalar inner product **d**^⊤^**1**_*N*_ is identical to 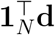, which we use throughout.

*Right eigenvector with eigenvalue* 0. We verify that **1**_*N*_ is the right null vector of **P**_*reg*_:

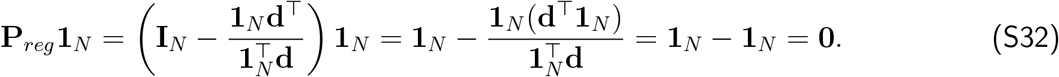

*Left eigenvector with eigenvalue* 0. We verify that **d**^⊤^ is the left null vector of **P**_*reg*_:

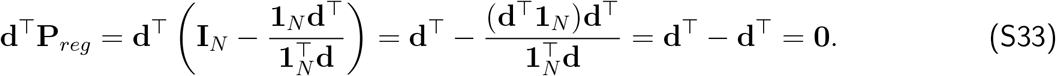

*Remaining eigenvectors with eigenvalue* 1. For any vector **v**_*k*_ satisfying **d**^⊤^**v**_*k*_ = 0:

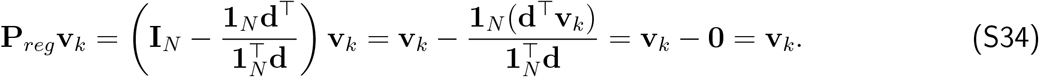

Thus **P**_*reg*_ has eigenvalues {0, 1, …, 1}, with right eigenvector **1**_*N*_ and left eigenvector **d**^⊤^ for eigenvalue 0, and all vectors in the null space of **d**^⊤^ as eigenvectors for eigenvalue 1.

### Proof of Proposition 3

We first show that Naive-GSR can be expressed as a spatial filter, then derive the post Naive-GSR covariance in terms of the degree vector.

Starting from the definition of the global signal 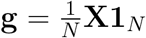:

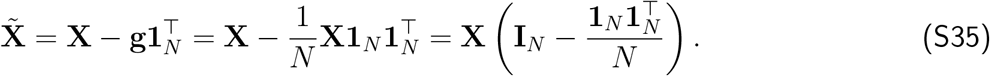

Thus 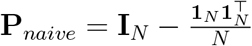.

The matrix **P**_*naive*_ is symmetric and idempotent 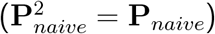, so it is an orthogonal projection. To find its eigenvalues:

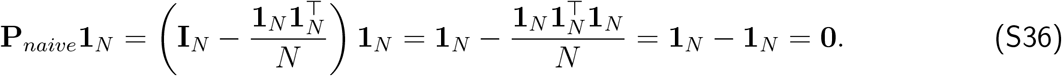

Thus **1**_*N*_ (or normalized 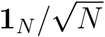) is an eigenvector with eigenvalue 0. For any vector **v** ⊥ **1**_*N*_ :

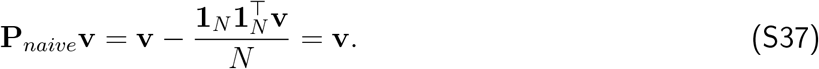

Thus all vectors orthogonal to **1**_*N*_ are eigenvectors with eigenvalue 1, giving eigenvalues {0, 1, …, 1}. The post Naive-GSR covariance is:

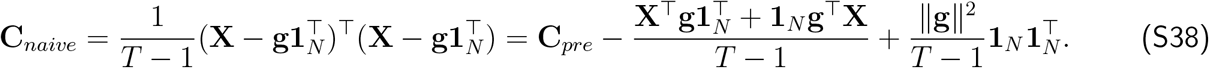

We express each term using the degree vector **d** = **C**_*pre*_**1**_*N*_ :

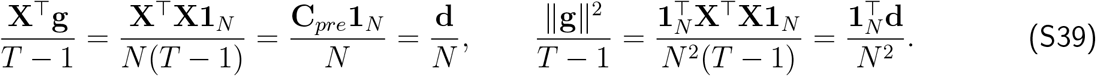

Substituting:

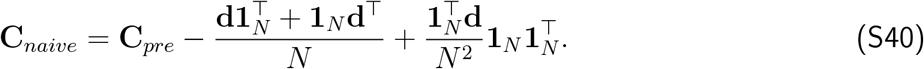

To verify that row sums are zero:

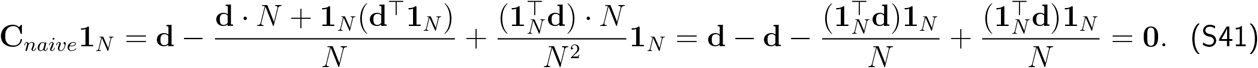

To establish the rank of the update, let Δ = **C**_*naive*_ − **C**_*pre*_. Every column of Δ is a linear combination of **d** and **1**_*N*_, so rank(Δ) ≤ 2. When 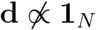, these vectors are linearly independent, giving rank(Δ) = 2. When **d** = *c***1**_*N*_ for some scalar *c*:

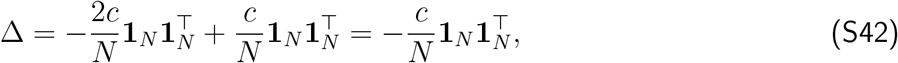

which is rank-1. This equals the Regression-GSR update 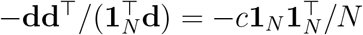, confirming that the two methods coincide when degree is uniform.

### Proof of Proposition 4

The definition 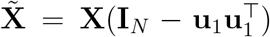 directly gives 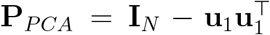. Since **u**_1_ is a unit vector 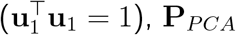 is symmetric and idempotent:

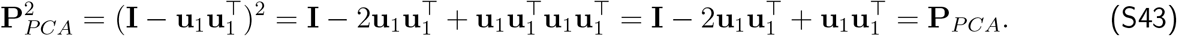

Thus **P**_*PCA*_ is an orthogonal projection onto the subspace orthogonal to **u**_1_.

The post PCA-GSR covariance can be written as:

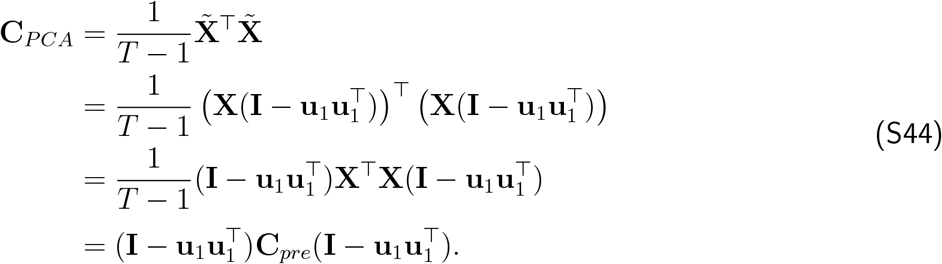

Expanding:

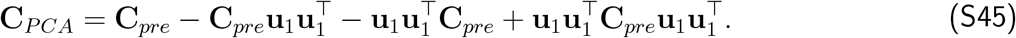

Since **C**_*pre*_**u**_1_ = *λ*_1_**u**_1_ and 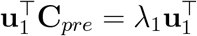:

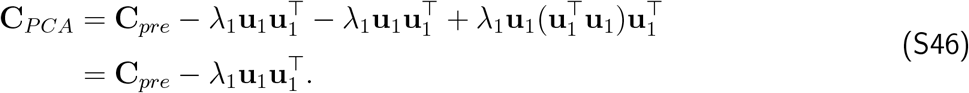

Using 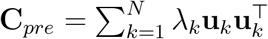:

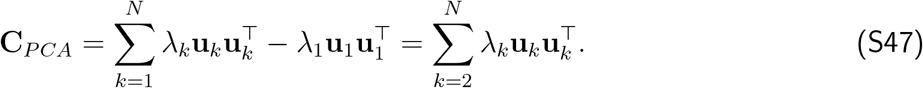

### Proof of Proposition 5

The symmetric normalized Laplacian is ℒ_*SC*_ = **I**−**D**^−1*/*2^**GD**^−1*/*2^. We verify that ***ψ***_1_ = **D**^1*/*2^**1**_*N*_ */*∥**D**^1*/*2^**1**_*N*_ ∥ is an eigenvector with eigenvalue 0:

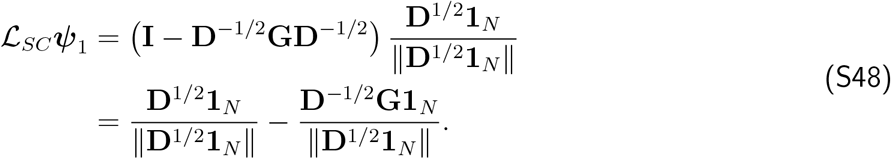

Since **G1**_*N*_ = **D1**_*N*_ (row sums of adjacency matrix equal degrees):

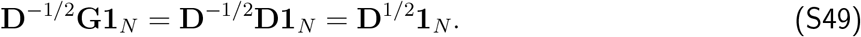

Therefore:

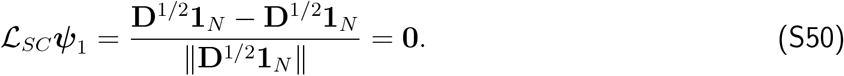

Thus ***ψ***_1_ is an eigenvector with eigenvalue 0.

The definition 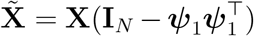 directly gives 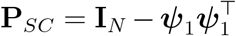, which is an orthogonal projection (symmetric and idempotent by the same argument as in Proposition 4).

## Footnotes

1 ^1^Data were provided by the Human Connectome Project, WU-Minn Consortium (Principal Investigators: David Van Essen and Kamil Ugurbil; 1U54MH091657) funded by the 16 NIH Institutes and Centers that support the NIH Blueprint for Neuroscience Research; and by the McDonnell Center for Systems Neuroscience at Washington University. Ethical approval was not required as confirmed by the license attached with the open access data.

